# Z-DNA as a Touchstone for Additive Empirical Force Fields and a Refinement of the Alpha/Gamma DNA torsions for AMBER

**DOI:** 10.1101/2021.07.11.451955

**Authors:** Marie Zgarbová, Jiří Šponer, Petr Jurečka

## Abstract

Although current AMBER force fields are relatively accurate for canonical B-DNA, many non-canonical structures are still described incorrectly. As non-canonical motifs are attracting increasing attention due to the role they play in living organisms, further improvement is desirable. Here, we have chosen Z-DNA molecule, can be considered a touchstone of the universality of empirical force fields, since the non-canonical α and γ backbone conformations native to Z-DNA are also found in protein-DNA complexes, i-motif DNA and other non-canonical DNAs. We show that spurious α/γ conformations occurring in simulations with current AMBER force fields, OL15 and bsc1, are largely due to inaccurate α/γ parameterization. Moreover, stabilization of native Z-DNA substates involving γ = trans conformations appears to be in conflict with the correct description of the canonical B-DNA structure. Because the balance of the native and spurious conformations is influenced by non-additive effects, this is a difficult case for an additive dihedral energy scheme such as AMBER. We propose new α/γ parameters, denoted OL21, and show that they improve the stability of native α/γ Z-DNA substates while keeping the canonical DNA description virtually unchanged, and thus represent a reasonable compromise within the additive force field framework. Although further extensive testing is needed, the new modification appears to be a promising step towards a more reliable description of non-canonical DNA motifs and provides the best performance for Z-DNA molecules among current AMBER force fields.

## Introduction

Force-field based modeling of nucleic acids is gaining popularity as a research tool, as ever-increasing computational power makes it possible to simulate ever larger systems and longer trajectories. However, better sampling in longer trajectories not only brings more possibilities, but also questions about the reliability and applicability of current nucleic acid force fields. On the one hand, we are increasingly confident about the canonical, or Watson-Crick double helical, structures of DNA and RNA, in simulations with the two most popular force field families, AMBER, with its well tested OL15^1^ and bsc1^2^ DNA variants and OL3^3^ for RNA, and CHARMM, with the non-polarizable force field CHARMM36^4^. It should be noted, however, that even for canonical DNA duplexes some unsolved issues remain, such as the poorly described A/B DNA equilibrium in the OL15 and bsc1 force fields.^5^ On the other hand, it is becoming increasingly apparent that there is a growing concern about non-canonical structures, where performance is arguably worse. In the case of the AMBER force fields, severe artifacts have been identified in RNA folding simulations,^6^ but inaccuracies have also been described in DNA force fields for guanine quadruplexes^7^ and for Z-DNA (although OL15 represents an improvement for Z-DNA, it is still not fully satisfactory, while bsc1 seems even further from the experimental Z-DNA conformation).^1,8^ This considerably limits the applicability of current force fields and calls for further work.

Non-canonical DNA motifs are receiving increasing attention as their relevance to the molecular biology of living organisms is being revealed. Unusual conformations are found in the contexts of DNA recombination (Holliday junction), viral DNA or telomere stability (DNA quadruplexes)^9^ and Z-DNA has been suggested to play a role in transcription.^10^ Also of interest is reliable description of non-canonical substates that may appear in unfolded DNA during its folding. Thus, the ability of our force fields to describe less common backbone conformations may be crucial in many applications.

### Z-DNA as a model molecule

In this work, we have chosen Z-DNA as our model molecule. Although a less common DNA form, it can be very useful for the force field development because its backbone is particularly sensitive to inaccuracies in the empirical potential. For example, the well known equilibrium of the ZI and ZII backbone substates in the GpC steps (Figure 1) has proven to be a difficult case for current force fields.^1,11^ Although this equilibrium has been much improved by the εζOL1 and βOL1 parameters (present in the OL15 force field, which is currently the best-performing force field for Z-DNA),^1,11^ there are other unusual backbone conformations in Z-DNA that are not modeled correctly by the current AMBER force fields, as discussed below. These unusual Z-DNA backbone substates involve non-canonical α = g+ and γ = t torsions, which are frequently found at protein-DNA interfaces (α/γ = g+/g-), in polycytosine i-motif structures, non-WC DNA pairs and DNAs intercalated with drugs (α/γ = g-/t).^12^ We would like to argue that this makes Z-DNA a very interesting test molecule that can serve as a simple and useful touchstone for the force field development, due to the number of coexisting backbone substates that model the variability of nucleic acids backbone in many other DNA structures.

**Figure 1.**
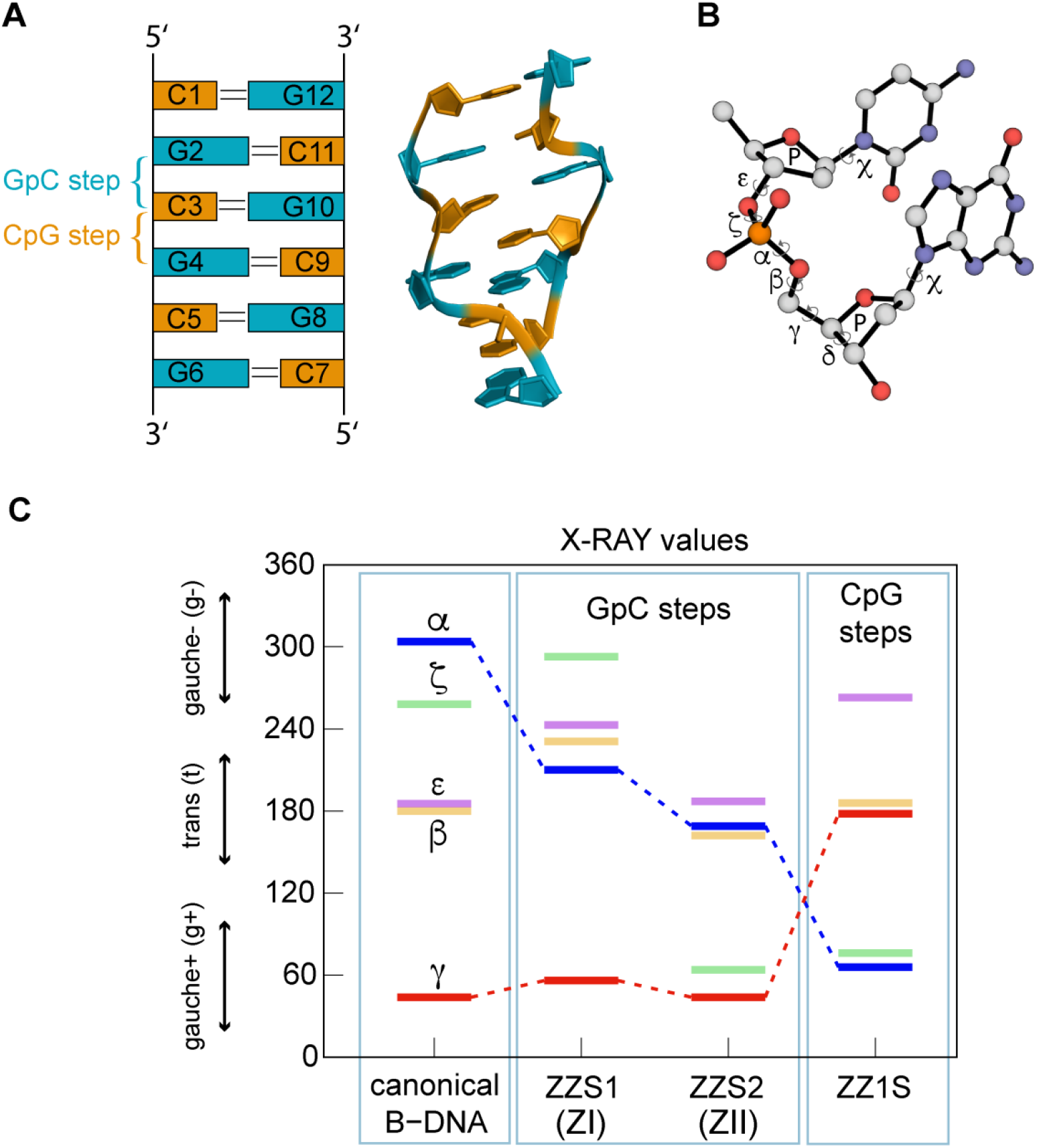
Z-DNA backbone substates. (A) GpC and CpG steps in Z-DNA, (B) backbone angle definition, (C) B-DNA (canonical) and Z-DNA (GpC and CpG steps) values of backbone dihedrals.

### Conflicting requirements for the description of B-DNA and Z-DNA

One conformation of the Z-DNA backbone that is difficult to model is the non-canonical α/γ = g+/t backbone substate that is native to the CpG steps of Z-DNA (Figure 1). As we will show below, this conformation is not sufficiently stable in the current AMBER force fields and transitions to the non-native α/γ = t/g+ conformation, which is a force field artifact (Table 1). When attempting to stabilize the α/γ = g+/t Z-DNA substate, it is important to realize that this could lead to deterioration of the B-DNA simulations, as the overpopulation of α/γ = g+/t in B-DNA was associated with a severe ff99 force field artifact that resulted in a large duplex undertwisting (tens of percent) on longer time scales (tens to hundreds of ns).^13^ This important historical artifact was corrected in 2007 by the bsc0 α/γ modification,^14^ which is also part of the bsc1 and OL15 force fields. Thus, the requirements for accurate description of B-DNA and Z-DNA are potentially contradictory, as stabilization of the native Z-DNA CpG step backbone (going against the bsc0 modification) may lead to deterioration of B-DNA double helix simulations, where the γ = t states are highly undesirable.

**Table 1.** Native and non-native α/γ backbone conformations in B-DNA and Z-DNA. Entangled system in which a conformation native to one system is an undesirable artifact in another.

| $\alpha/\gamma$ conformation | $g-/g+$ | $g+/t$ | $t/g+$ |
| --- | --- | --- | --- |
| B-DNA | native | FF artifact | - |
| Z-DNA, GpC steps | - | - | native (ZI, ZII) |
| Z-DNA, CpG steps | - | native | FF artifact |

Alternatively, instead of stabilizing the native Z-DNA CpG backbone, destabilization of the non-native α/γ = t/g+ conformation can be considered to suppress its occurrence. However, since this conformation is native in the ZI/ZII conformations of Z-DNA GpC steps (Table 1), its destabilization could negatively affect their modeling. Note that instability of the GpC steps, specifically the occurrence of some non-native substates, was observed in our earlier work.^1^ It is clear that a balanced description of all substates involved (native or spurious) is necessary for accurate modeling of both B-DNA and Z-DNA. We would like to note that achieving a satisfactory balance may not even be possible due to the non-additivity of some backbone dihedral angle energies, namely α/ζ, which cannot be modeled within the additive dihedral scheme of AMBER (see below). Thus, it remains a challenge to find out to what extent AMBER force field form is able to accurately model the equilibria in Z-DNA.

There are several possible reasons for the poor modeling of Z-DNA backbone by current force fields. One of them is the coupling with neighboring dihedrals through hyperconjugation (or anomeric) effect extending through the phosphate group.^15^ Meaningful description of this effect would require non-additive model of the neighboring torsions that could be (partially) achieved in non-additive force fields^16–17^ or by applying a CMAP-type correction as in CHARMM.^18–19^ In what follows, however, we explore the possibility that the current parameterization could be improved even in the additive torsion framework.

Another straightforward reason for poor Z-DNA modeling could simply be the inaccuracy of α/γ dihedral parameterization. While the ff94^20^ and ff99^21^ force fields used the original ff94 α/γ parameters, the two currently most commonly used force fields for DNA, bsc1^2^ and OL15^1^ use the bsc0 α/γ dihedral refinement by Perez et al.^14^ This refinement was also recommended for RNA simulations to support the essential χ_OL3_ refinement.^22^ For the sake of completeness, other RNA-specific reparameterizations involving α/γ emerged recently.^23–25^ Based on simulations of various RNA molecules, we have previously hypothesized that the γ = t region may be overly destabilized in the bsc0 refinement. Indeed, the poor stability of the α/γ = g+/t substate in Z-DNA discussed here is consistent with this notion. Therefore, we will try to find α/γ refinement that would improve the modeling of non-canonical substates native in Z-DNA while not affecting the good performance of OL15 force field for the canonical B-DNA.

To derive a new α/γ parameterization, we used fitting to multiple QM reference curves. A unique feature of our methodology is that it includes conformation-dependent solvation effects while avoiding double counting of solvation energy.^26^ However, as discussed below, the resulting fits were afterwards modified in some regions to better balance the description of the various important backbone conformations.

In our previous work we have noted that water model used may affect the subtle equilibrium of Z-DNA substates. When the SPC/E water model is used instead of TIP3P, population of the non-native α/γ = g-/g+ state in the CpG steps of Z-DNA, which is central to this work, decreases from 42% to 22% in ff99bsc0βol1εζol1χol4 (i.e., OL15) simulation.^1^ Therefore, we will consider here three different water models, SPC/E, TIP3P and OPC. In addition, in 2012 Steinbrecher, Latzer and Case published a modification of the phosphate vdW parameters that, among other effects, improved the solvation energy of the phosphate group.^27^ This modification, hereafter referred to as SLC, proved to be a partial improvement in RNA simulations, where it helped to reduce the populations of misfolded structures in folding simulations.^6,28^ Because these parameters appear to be more physically plausible than the original parm99 phosphate, we decided to test them here as well.

In this paper, we start by describing the α/γ modification and its effect on the α/γ Z-DNA backbone. Then we investigate the effect of water and SLC phosphate parameters on Z-DNA. Finally, we evaluate the effect of the new parameterization on a canonical B-DNA duplex, in particular the population of non-canonical α/γ backbone substates and the possible effect on a range of its structural parameters.

## Methods

### Derivation of α/γ parameters

In deriving the new α/γ parameters, we used a previously described method that includes conformation-dependent solvation effects while avoiding double counting of solvation energy.^26^ The same method was used to derive the previous “OL” adjustments (χ_OL3_, χ_OL4_, εζ_OL1_, β_OL1_).^1,3,7,11^ All calculations were performed on a small model compound (Figure 2). In order to cover most important α/γ combinations a total of four energy profiles were considered, two for the αscan with γconstrained to *gauche+* (50°) and *trans* (180°), and two for the γ scan with α constrained to *gauche+* (60°) and *gauche*- (300°); all remaining inner coordinates were relaxed (starting from the canonical values) and scanning was performed in 10° steps.

**Figure 2.**
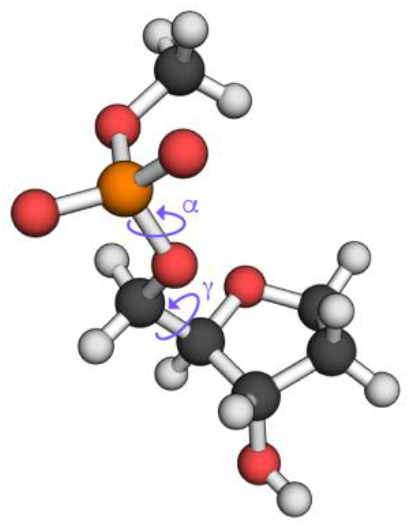
Model compound for α/γ QM scans.

The QM reference curves were obtained using high-level QM calculations and include specific solvation effects. First, all geometries were optimized at the PBE/QZVPP level with the COSMO solvent model^29^ and ε_r_ = 78.4. The default scaled Bondi radii with the scaling factor 1.17 were used. Then, high-level single-point QM calculations were performed on the optimized geometries using the RI-MP2/CBS +ΔCCSD(T) calculations. The complete basis set (CBS) extrapolations were realized through the scheme of Helgaker and Halkier^30–31^ (HF and MP2 energies were extrapolated separately) using aug-cc-pVTZ and aug-cc-pVQZ basis sets. The correction term for higher-order correlation effects, ΔCCSD(T), was calculated using the aug-cc-pVDZ basis set. All QM calculations were performed using Turbomole 6.3.^32^ Molecular mechanical optimizations were performed in Gaussian03^33^ using the “external” function and an inhouse script linking Gaussian to the sander module of AMBER 9.^34^ In force field calculations, the default charges were used for all atoms except the hydrogens on the terminal methyl group and carbon C1’, where they were set so that the total charge of the compound was −1. Poisson-Boltzmann (PB)^35–36^ model was used as a solvent model with the grid spacing of 0.1 Å, ε_r_ =78.4 and the same radii as in QM [see Ref. ^26^]. The resulting parameters are given in Supplementary Information, Table S1.

### MD simulations

Table 2 details force field combinations used in this work. The newly derived parameters are denoted by OL21 α/γ or simply OL21 in the following, where “OL” stands for the city of Olomouc and in our work they are compared with two currently most widely used DNA force fields, bsc1^2^ and OL15.^1^

**Table 2.** An overview of the DNA dihedral parameters used.

| Combination | parent FF | $\alpha/\gamma$ | $\beta$ | $\epsilon/\zeta$ | $\chi$ | Pucker |
| --- | --- | --- | --- | --- | --- | --- |
| OL15 | parm99<sup>21</sup> | bsc0<sup>14</sup> | $\beta_{OL1}^1$ | $\epsilon/\zeta_{OL1}^{11}$ | $\chi_{OL4}^7$ | parm99 |
| OL21 | parm99 | OL21 $\alpha/\gamma$ | $\beta_{OL1}^1$ | $\epsilon/\zeta_{OL1}^{11}$ | $\chi_{OL4}^7$ | parm99 |
| bsc1 | parm99 | bsc0 | parm99 | bsc1 $\epsilon/\zeta^2$ | bsc1 $\chi$ | bsc1 P |

The starting structures were d(CGCGCG)_2_ duplex (PDB ID 1ICK^37^; X-ray resolution 0.95 Å; spermine and magnesium removed) for Z-DNA and the Drew-Dickerson dodecamer d(CGCGAATTCGCG)_2_ (PDB ID 1BNA^38^; X-ray resolution 1.9 Å) for B-DNA. An overview of the simulations and simulation conditions is given in Table 3.

**Table 3.** An overview of MD simulations.

| Molecule | Start PDB ID | Force field | Salt | Water model | Time [μs] | Duplex ends | Phosphate vdW |
| --- | --- | --- | --- | --- | --- | --- | --- |
| B-DNA | 1BNA | OL15<br>OL21<br>bsc1 | 0.15M KCl | SPCE | 2 | free | original |
| B-DNA | 1BNA | OL15<br>OL21<br>bsc1 | 0.15M KCl | SPCE | 2 | restrained<sup>a)</sup> | original |
| Z-DNA | 1ICK | OL15<br>OL21<br>bsc1 | 2M NaCl | SPCE<sup>39</sup> | 5 | free | original |
| Z-DNA | 1ICK | OL15<br>OL21<br>bsc1 | 2M NaCl | SPCE | 5 | free | SLC |
| Z-DNA | 1ICK | OL21 | 2M NaCl | OPC<sup>40</sup> | 5 | free | original |
| Z-DNA | 1ICK | OL21 | 2M NaCl | TIP3P<sup>41</sup> | 5 | free | original |

<sup>a)</sup> Flat well restraint between the electronegative atoms of terminal base pair hydrogen bonds, flat between 2.5-3.2 Å, parabolic restraints with $k = 20 \text{ kcal}/(\text{mol} \cdot \text{Å}^2)$ for shorter and $30 \text{ kcal}/(\text{mol} \cdot \text{Å}^2)$ for longer distances, respectively.

The initial structures were first neutralized with Na^+^ (Z-DNA) or K^+^ (B-DNA) cations, and the ion-concentration was then adjusted to 2M-NaCl (Z-DNA) or 0.15M-KCl (B-DNA) using ionic parameters by Joung and Cheatham.^42–43^ In selected Z-DNA simulations, alternative vdW parameters for phosphate oxygen (SLC, see above) were used.^27^ An octahedral box with a 12 Å buffer was used to solvate the molecular systems. Molecular dynamics simulations were performed using the PMEMD code (for CUDA) from the AMBER 16 and 18^34^ program suits under NPT conditions (1bar, 298 K) with the Langevin thermostat temperature control, Monte Carlo barostat pressure control (taup=2), a 4 fs time step (using hydrogen mass repartitioning procedure^44–45^), a 10 Å direct space nonbonded cutoff, and SHAKE on bonds on hydrogen atoms with default tolerance (0.00001 Å). The nonbonded pair list was updated every 25 steps. Coordinates of nucleic acids and ions were stored every 10 ps.

In order to compare the B-DNA helical parameters unaffected by the stochastic fraying effects, we performed another set of Drew-Dickerson duplex simulations in which flat well distance restraints were applied between the electronegative atoms of terminal base pair hydrogen bonds (“restrained” in Table 3; see footnote).

The analysis of nucleic acid structure parameters was performed using the cpptraj module and nastruct tool of AMBER software package. The calculation of the groove widths was performed following the approach of El Hassan and Calladine.^46^

### Classification of backbone conformations

We use the nomenclature of DNA/RNA dinucleotide conformations according to Černý et al.^47^ Substates ZI(ZZS1)/ZII(ZZS2) and ZZ1S are assigned based on the shortest distance in the nine-dimensional space of the dihedral angles of two consecutive nucleotides 1 and 2 (δ1, χ1, ε1, ζ1, α2, β2, γ2, δ2 and χ2) from the representative X-ray values by Černý et al.^47^ This method is simpler than the method proposed in the original work, which was based on a machine learning technique, but it provides satisfactory results for Z-DNA. If the distance to the closest conformer (ZI, ZII for GpC step or ZZ1S for the CpG step) in the 9D dihedral space is greater than 80 degrees cutoff, the conformer is unassigned (denoted “other”). Note that in our previous work^1^ we assigned conformers based on whether they belonged to a certain dihedral angles interval, not based on the shortest distance criterion described above, so the results may vary slightly.

## Results and Discussion

### Parameter development

We started our derivation by fitting reference (QM – MM) curves which include conformation-dependent solvation effects, described as the difference between the QM solvation energy (naturally including solute polarization effects) and MM solvation, in which the conformation-dependent solute polarization is neglected.^26^ Thus, the derived parameters do not include full solvation energy, only the mentioned difference. Note however, that the reference curves shown in Figure 3 include the full solvation energy, both for QM and MM calculations. The newly derived parameters are denoted by OL21 α/γ or simply OL21 in the following and are shown in Figure 3 (torsion potentials) and Figure 4 (dihedral parameters).

**Figure 3.**
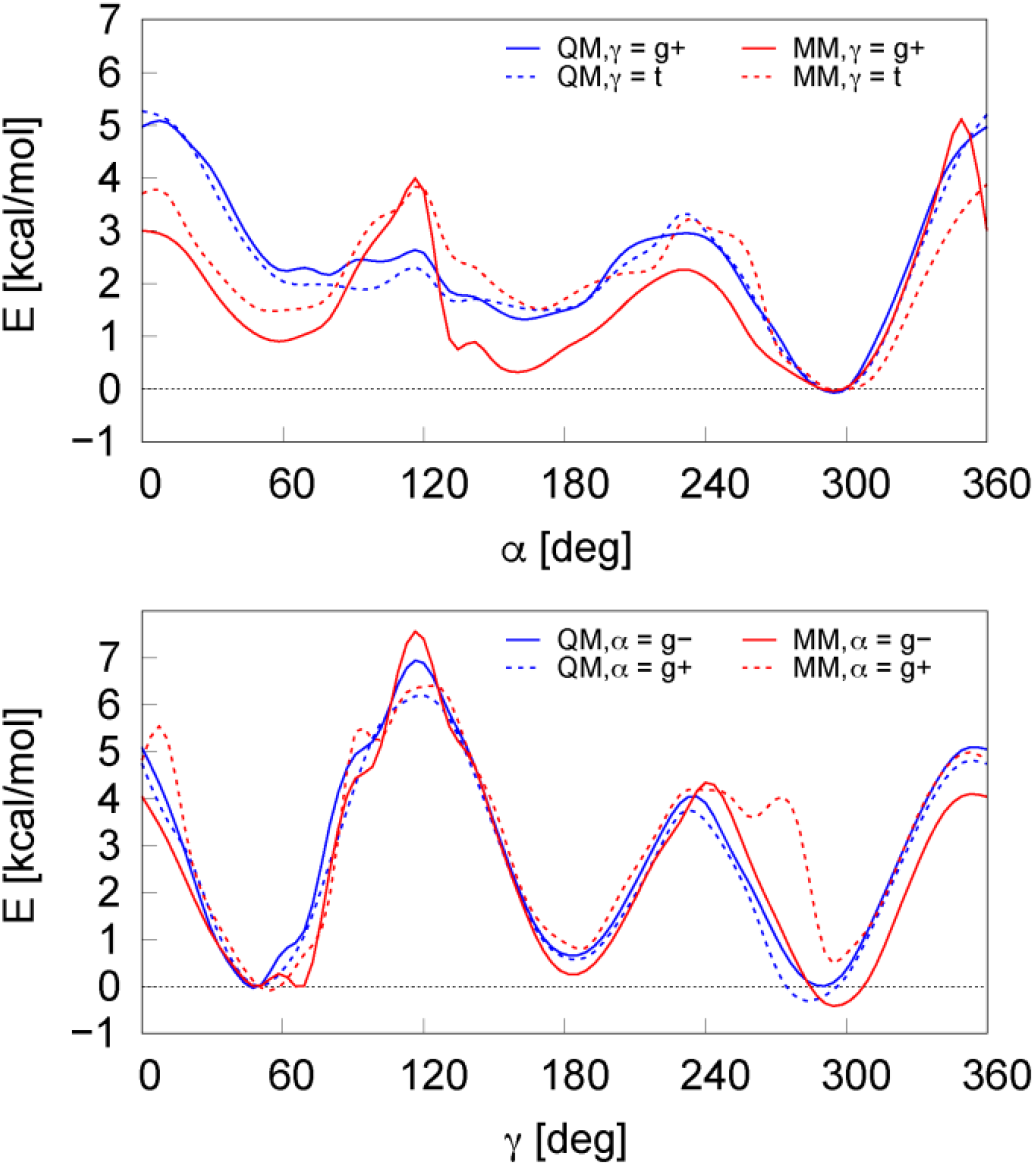
QM scans and OL21 fits for the α torsion (with γ set to g+ and t) and the γ torsion (with α set to g+ and g-).

**Figure 4.**
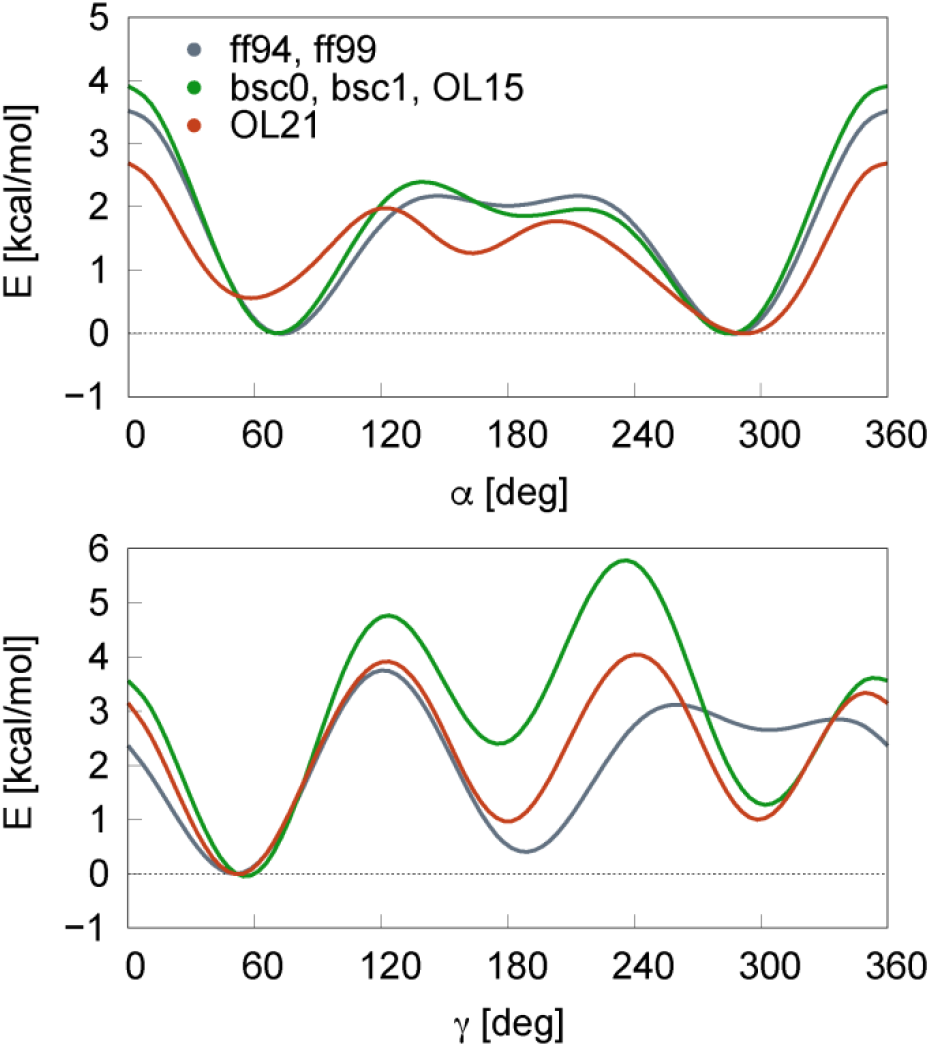
Dihedral term potential energy for OL21 α/γ compared with previous parameterizations.

The resulting α profiles are a compromise between reference scans for two different fixed γ angles (and vice versa for the γ angle). We would like to note that the OL21 α/γ parameters are not purely a fit to the reference data, but are in part a carefully tuned practical compromise allowing a balanced description of various backbone substates. This mainly concerns the γ = t region, which was found to be excessively destabilized by the original (bsc0) force field in our previous work,^48^ and also the α = g+ region, which is tightly connected to ζ dihedral through hyperconjugation and may not be fully satisfactorily described in additive dihedral scheme such as AMBER (see above).

The differences with respect to the previous parameterizations are best seen in Figure 4. The α = t and γ = t regions have been changed to approximately match the α/γ = t/t conformation energy relative to the canonical α/γ = g+/g-. The resulting γ = t potential minimum is now much lower than in the original bsc0 parameterization (see Figure 4), but still in reasonable agreement with our reference QM data. This could help to reduce the aforementioned over-destabilization of the γ = t in bsc0^48^ and thus help stabilize the native ZZ1S in CpG step of Z-DNA, which is of our interest in this work. However, we will have to carefully monitor B-DNA simulations for an excessive population of the spurious α/γ = g+/t state, which was a major problem of the ff99 force field.^13^

The α = g+ region was destabilized with respect to previous parameters (Figure 4), but less than indicated by the reference QM data (Figure 3). The destabilization of α = g+ region in the QM scan is due to the unfavorable ζ/α = g-/g+ conformation, which is energetically higher than the ζ/α = g+/g+, g-/g- and g/t combinations.^16–17^ The partial destabilization in our potential reflects a compromise between possible ζ/α combinations and the quality of this compromise will be reflected in our MD simulations, particularly in the populations of the undesired (spurious) α/γ = g+/t substates in B-DNA simulations vs. the stability of the ZZ1S substate in the CpG steps of Z-DNA.

### Z-DNA Simulations and Backbone Conformation

The overall fold of Z-DNA is well preserved by the OL15 and OL21 force fields, which provide a relatively small RMSD from the X-ray structure within the 5 μs of our simulation (Figure 5), consistent with low RMSD observed before for the bsc0 force field.^1^ The bsc1 simulation shows much larger RMSD fluctuations. The first 400 ns are similar to those reported in the 400 ns simulation in Figure 2a of the original paper,^2^ but the later stages of the bsc1 simulation show a major increase of RMSD, consistent with previous studies.^8^ We find that most of the bsc1 RMSD deviation is due to the instability of the terminal Z-DNA base pairs, with the terminal bases spending some time free in solution, but most of the time hydrogen bonded in groove of the Z-DNA helix by various hydrogen bonds involving guanine -NH_2_ group. The resulting structures often feature a 5’-OH group hydrogen bonded to a carbonyl oxygen of the inner cytosine, and we have previously suggested that this bond may be overestimated in current force fields and may be a main source of non-native frayed structures in B-DNA helices.^49^ Nevertheless, the RMSD of the bsc1 simulation is notably higher even before fraying occurs and in the inner stem region throughout the simulation (not shown). This is mainly due to the non-native (spurious) α/γ = t/g+ backbone substates, which are heavily populated in bsc1 Z-DNA simulation (which we will discuss in more detail below).

**Figure 5.**
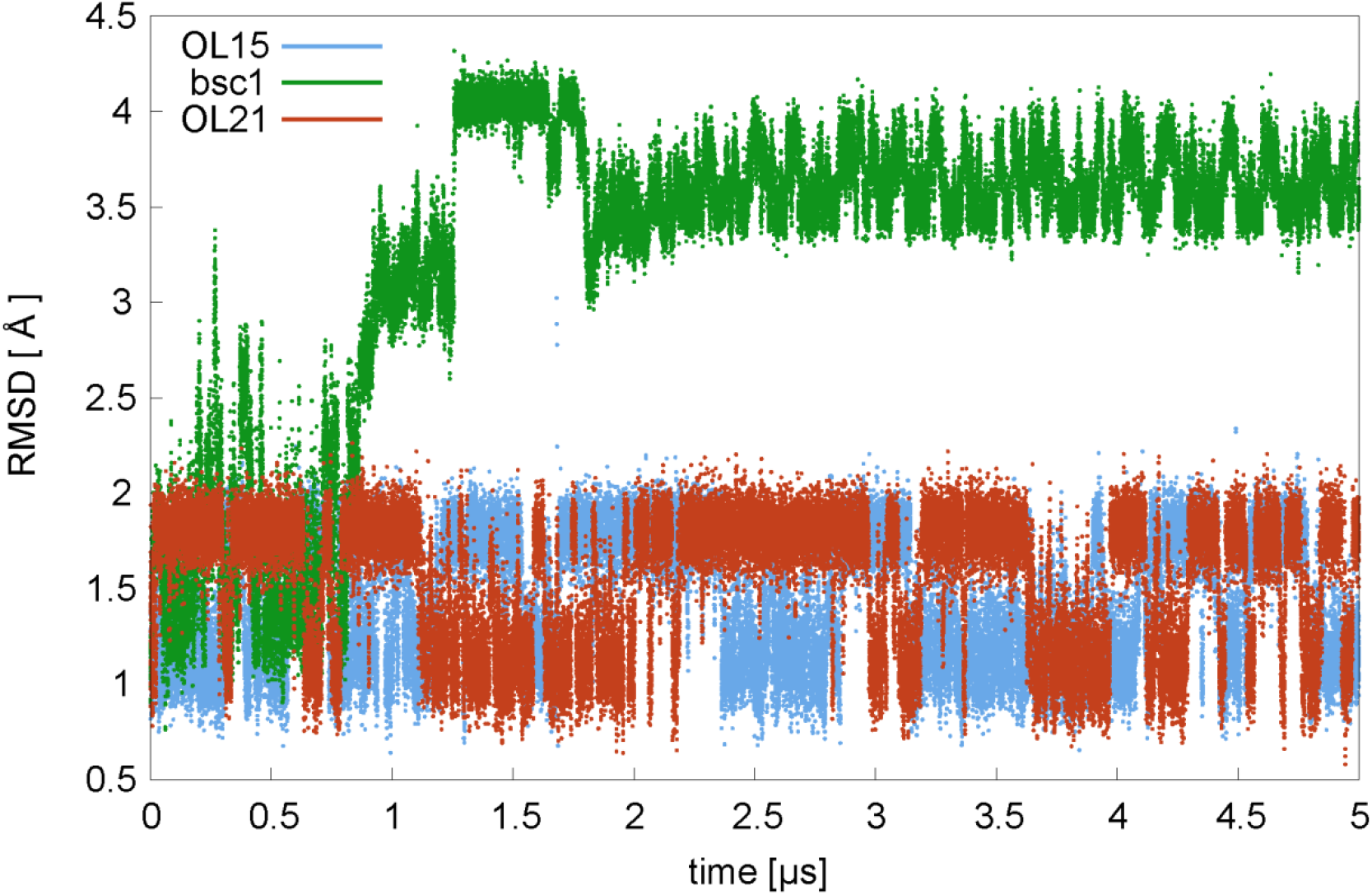
Z-DNA RMSD for OL21, bsc1 and OL15 simulations in SPC/E water with original phosphate vdW parameters.

We can now take a closer look at the backbone dihedrals that define the Z-DNA conformational substates, separately for GpC and CpG steps (Figure 6). An overview of native and selected non-native (spurious) substates is given in Table 4. Let us start with the GpC steps, for which the ZI/ZII equilibrium discussed in our earlier work for OL15^1^ is only slightly affected by the addition of the OL21 α/γ parameters to OL15 (right panel). The ZI conformation is dominant (90%, see Table 5), ZII is minor (7%) and the admixture of other (non-native) substates remains low for the simulations with the original AMBER phosphate parameters. The results are similar when using SLC phosphate vdW parameters instead of the original parameters. In contrast, the bsc1 simulation shows only 18% of the native ZI and 1% of the ZII substate, and the remaining snapshots differ markedly from the reference X-ray data. To see what is happening in the bsc1 simulation, we can look at Figure 6. While the distributions of the ζ, α and γ angles are very similar to the OL15 distributions for the ZI conformation, those of the ε and β torsions are closer to the ZII conformation. Along with this, the δangle on guanine shifts from values typical for the N conformation to the S conformation, and the χ angle shifts from the *syn* region (around 60°) to around 120°. Thus, bsc1 models a state that is inconsistent with the X-ray reference. We speculate that this could be due to the difference in the β potential between bsc1 and OL15 – the OL15 β derived from the QM fit has a lower energy penalty for the β^~^ 230° region, which is characteristic of the native ZI conformation. The QM fit of OL15 therefore describes this region better than the original ff99 AMBER β potential that is used also in the bsc1 force field. Note that the Z-DNA β distributions similar to that of bsc1 have also been reported for earlier versions of the force field before the β_OL1_ refinement was introduced.^1^

**Figure 6.**
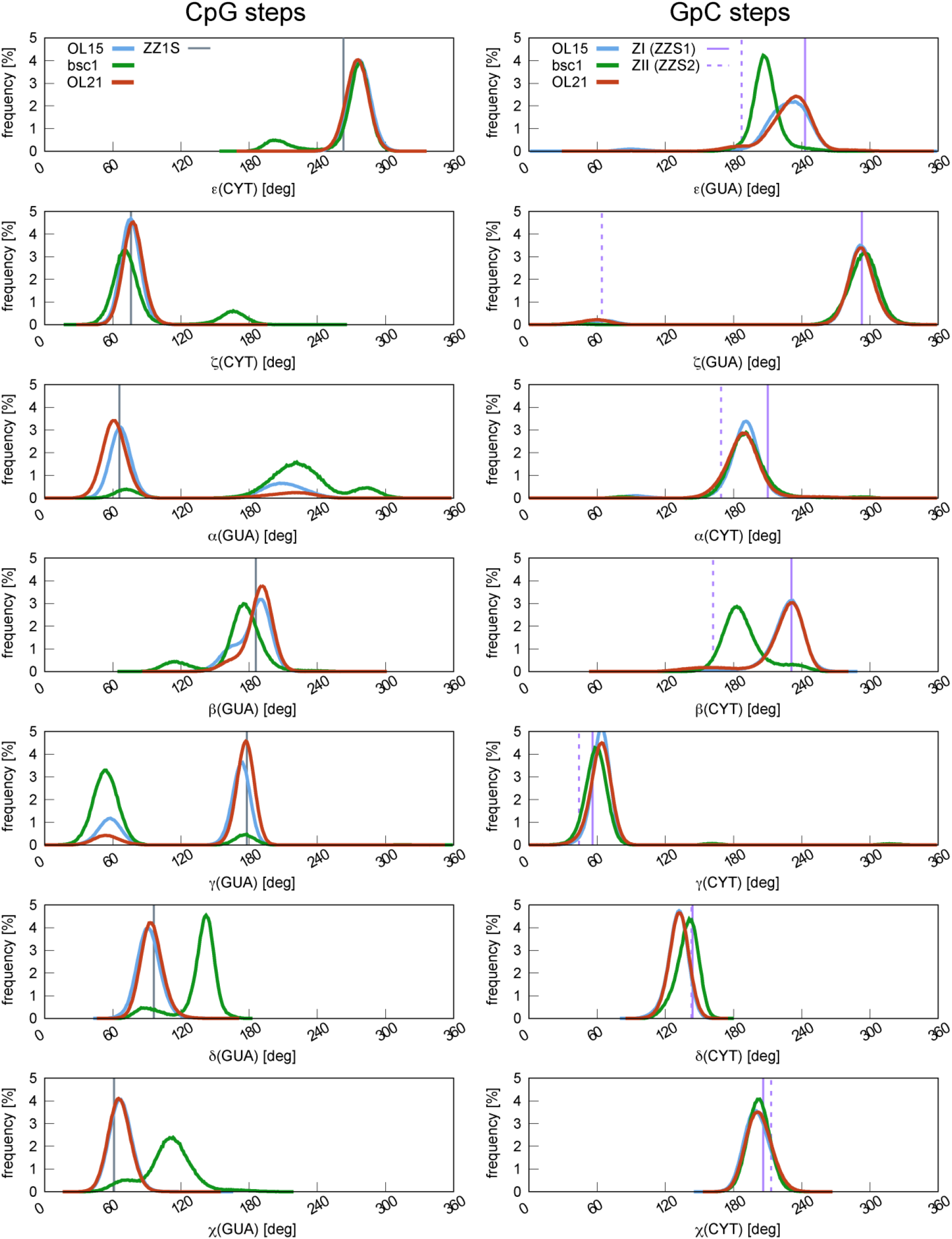
Dihedral angle distributions in CpG and GpC steps of Z-DNA^a)^. Reference X-ray values^47^ are indicated by vertical lines for ZI (ZZS1, solid line) and ZII (ZZS2, dashed line) and by vertical solid lines for the native α/γ = g+/t (ZZ1S) conformation. Original phosphate vdW parameters and SPC/E water, 5 μs. ^a)^ Distributions are shown for inner and 5’-end steps only; 3’-end steps are excluded because they are in ZZ2S conformation. For the bsc1 simulation, only the part of the simulation that did not exhibit fraying was included to avoid contamination due to fraying.

**Table 4.** Canonical B-DNA (BI) backbone dihedrals compared with native GpC ZI/ZII and CpG ZZ1S conformations of Z-DNA and spurious non-native substates in CpG steps.

| Backbone substate | Dihedral angle and sugar pucker P |  |  |  |  |  |  |  |  |
| --- | --- | --- | --- | --- | --- | --- | --- | --- | --- |
| | $\varepsilon$<br>(C) | $\zeta$<br>(C) | $\alpha+1$<br>(G) | $\beta+1$<br>(G) | $\gamma+1$<br>(G) | P<br>(C) | P+1<br>(C) | $\chi$<br>(C) | $\chi+1$<br>(G) |
| B-DNA canonical | t | g- | g- | t | g+ | S | S | high-<br>*anti* | high-<br>*anti* |
| GpC steps |  |  |  |  |  |  |  |  |  |
| ZI (ZZS1<sup>a)</sup>) | g- | g- | t | ~g- | g+ | N | S | *syn* | *anti* |
| ZII (ZZS2<sup>a)</sup>) | t | g+ | t | t | g+ | N | S | *syn* | *anti* |
| CpG steps |  |  |  |  |  |  |  |  |  |
| native (ZZ1S<sup>a)</sup>) | g- | g+ | g+ | t | t | S | N | *anti* | *syn* |
| non-native OL21 | g- | g+ | t | t | g+ | S, 130° | N | *anti* | *syn* |
| non-native bsc1 | t | g- | t | t | g+ | S, 150° | S | *anti* | 120° |

<sup>a)</sup> See Ref. <sup>47</sup>

**Table 5.** Populations of ZI/ZII and α/γ = g+/t (ZZ1S, inner + 5’end) substates and the effect of phosphate vdW parameters (original vs. SLC). SPC/E water, 5 μs.

| SPC/E | % of the native substate in GpC (ZI/ZII) and CpG (ZZ1S) steps |  |  |  |  |  |
| --- | --- | --- | --- | --- | --- | --- |
|  | Original phosphate vdW |  |  | SLC phosphate vdW |  |  |
|  | OL15 | bsc1<sup>a)</sup> | OL21 | OL15 | bsc1<sup>a)</sup> | OL21 |
| GpC steps |  |  |  |  |  |  |
| ZI (ZZS1) | 92 | 18 | 90 | 92 | 42 | 87 |
| ZII (ZZS2) | 3 | 1 | 7 | 5 | 7 | 10 |
| sum native | 95 | 19 | 97 | 97 | 49 | 97 |
| CpG steps |  |  |  |  |  |  |
| ZZ1S native | 69 | 6 | 86 | 80 | 32 | 92 |

<sup>a)</sup> For the bsc1 simulation, only the part of the simulation that did not exhibit fraying was included to avoid contamination due to fraying.

We now turn to the main interest of this work, the α/γ backbone conformation in the CpG step (left column in Figure 6). Major deviations from the X-ray reference are observed for the bsc1 force field, where only 6% of the trajectory corresponds to the native substate ZZ1S (α/γ = g+/t). Instead, a nonnative distribution with α^~^ 180-300° and γ = g+ is observed, a force field artifact inconsistent with the X-ray structures of Z-DNA. With OL15, population of the native ZZ1S is 69%, which is much better, but still not entirely satisfactory. The newly derived OL21 with the γ = t region lowered relative to OL15 and bsc1 provides stabilization of the native ZZ1S state and better agreement with the experiment (86% of the native population with the original AMBER phosphate vdW parameters and 92% with the SLC parameterization).

### Z-DNA backbone is sensitive to water model and phosphate vdW parameters

In our earlier work we noted that the water model has an important effect on the Z-DNA backbone substates. When SPC/E water was used instead of TIP3P, the non-native α/γ = t/g+ state population of in the CpG step decreased from 42% to 22% in ff99bsc0βol1εζol1χol4 (OL15) simulation.^1^ Here we studied three water models, TIP3P, SPC/E and OPC with OL21 force field, both with the original and Steinbrecher-Latzer-Case (SLC) phosphate parameters (Table 6). Again, we see that the TIP3P model reduces the population of native Z-DNA substates in the CpG step, but to a lesser extent than with the OL15 force field. In addition, we can see that the OPC model also appears to stabilize the non-native substate, although to a lesser extent than TIP3P. Similar effects can also be observed for the ZI/ZII equilibrium in the GpC steps, with SPC/E water model again providing the best description in terms of the largest populations of the native (ZI, ZII) backbone conformations.

**Table 6.** Effect of water model and phosphate vdW parameters on ZI/ZII and α/γ = g+/t (ZZ1S, inner and 5’ends) substates in 5 μs OL21 simulation.

| OL21 | % of the native substate in GpC (ZI/ZII) and CpG (ZZ1S) steps |  |  |  |  |  |
| --- | --- | --- | --- | --- | --- | --- |
|  | Original phosphate vdW |  |  | SLC phosphate vdW |  |  |
|  | TIP3P | SPC/E | OPC | TIP3P | SPC/E | OPC |
| GpC steps |  |  |  |  |  |  |
| ZI (ZZS1) | 85 | 90 | 82 | 84 | 87 | 80 |
| ZII (ZZS2) | 7 | 7 | 10 | 10 | 10 | 13 |
| sum | 92 | 97 | 92 | 94 | 97 | 93 |
| CpG steps |  |  |  |  |  |  |
| ZZ1S (CpG steps) | 80 | 86 | 83 | 86 | 92 | 90 |

The SLC phosphate parameters also appear to affect both the ZI/ZII and ZZ1S substates. The strongest effect is seen for the bsc1 force field, where SLC parameters increase the percentage of native ZI conformation from 18% to 42% and native ZZ1S conformation from 6% to 32% (Table 5). A similar positive effect is also seen for the OL15 and OL21 force fields, although to a lesser extent. A small stabilization of the ZII substate relative to ZI by SLC phosphates is also desirable. In conclusion, SLC phosphates seem to provide a somewhat better description of Z-DNA than the original phosphate parameterization and can be recommended as viable alternative parameters. Nevertheless, the improvement achieved by the OL21 α/γ refinement is more important than the effect of the phosphate vdW parameters.

### B-DNA remains virtually unchanged by the OL21 α/γ modification

Because we did not significantly modify the α/γ potentials near their canonical minima, we did not expect any significant changes in canonical B-DNA structure and dynamics. Indeed, we found that the most studied DNA duplex, Dickerson-Drew dodecamer (DDD), is almost unaffected by the OL21 α/γ modification. This can be seen from the base pair step helical parameters and their sequence dependence for which deviations from the OL15 force field were very small (Figure S1). Also, the histograms of the backbone angles remained virtually unchanged (Figure S2) as well as RMSD (Figure S3; average RMSD was 1.25, 1.32 add 1.26 Å for OL15, bsc1 and OL21, respectively). The total percentage of BI/BII conformations changed only slightly (24.5 % with OL15 vs. 24.7 % with OL21) and its sequence dependence is again very similar (Figure S4). Averages of selected geometrical parameters of Dickerson-Drew dodecamer are given in Supporting Information, Table S2.

A major concern when modifying the γ potential in OL21 was the stabilization of the γ= t region. As mentioned in the Introduction, we had to avoid over-stabilization of the γ = t region that existed in ff99 and led to the accumulation of the α/γ = g+/t backbone substates, a severe force field artifact that was corrected in 2007 by the bsc0 α/γ modification,^14^ which has been transferred to the bsc1 and OL15 force fields. Therefore, the population of unwanted α/γ substates is clearly an important issue. Table 7 shows the populations of non-native α/γbackbone conformations in our 2 μs simulation. The population of the mentioned α/γ = g+/t in the OL21 simulation is very small, approximately 1 %, which should have no significant effect on B-DNA properties. The only other conformations detected are 0.3 % of α/γ = t/t and 0.1 % of α/γ = g+/g-, which are negligible. Time series of α, γ, and β dihedral angles showing the frequency of non-canonical backbones for all three force fields are given in Supporting Information, Figures S5-S7. We would like to note that small populations of non-canonical α/γ backbone substates in DNA simulations are probably desirable, as some of them may play a role in DNA conformational landscape, for example in stabilizing protein-DNA complexes.^12^ Note that the α/γ = g+/t geometry is the one found in the CpG steps of Z-DNA, where it is very stable in our OL21 simulations. This shows that our α/γ parameters represent a very good compromise between the stability of native non-canonical α/γ Z-DNA (a representative of non-canonical DNA structures) backbone conformers and canonical B-DNA.

**Table 7.** Populations (%) of α/γ backbone substates in 2 μs B-DNA DDD simulation with SPC/E water model for OL15, bsc1 and OL21 force fields.^a)^

| Backbone conformation |  | OL15 | bsc1 | OL21 |
| --- | --- | --- | --- | --- |
| $\alpha/\gamma$ | g-/g+ (canonical) | 99.3 | 97.2 | 98.5 |
|  | g+/t (FF artifact<sup>13</sup>) | 0.3 | 0.6 | 0.9 |
|  | t/t | 0.0 | 0.2 | 0.3 |
|  | g+/g- | 0.0 | 0.0 | 0.1 |
| $\beta/\gamma$ | t/g+ | 0.0 | 2.0 | 0.0 |

<sup>a)</sup> Simulation in 0.15 M KCl and SPC/E water, with original phosphate vdW parameters and flat well restraints at both ends to avoid bias from occasional fraying.

Fraying of the ends of DNA double helix is another noteworthy issue. Figure S8 of the Supporting Information shows fraying (a consequence of terminal base pair breaking) for all three studied force fields. All three force fields show some degree of fraying at one or both ends of the helix. Only the terminal base pair is involved and the structures formed are those described in our previous work on fraying of DNA and RNA [ref.^49^] Since the nature of the fraying events is stochastic and they occur on relatively long time scales, we cannot say whether any of the force fields are clearly different or better on the current simulation time scale.

## Conclusions

The Z-DNA molecule contains non-canonical α and γ backbone conformations, which are of wider interest due to their occurrence in protein-DNA complexes, i-motif DNA and other non-canonical and unfolded DNAs. One such conformation is the ZZ1S conformation with the α/γ = g+/t combination, which, however, is poorly modeled by current AMBER force fields. Moreover, the α/γ = g+/t substate stabilization required for Z-DNA may lead to its overpopulation in the B-DNA double helix, which has previously been shown to be a major problem. The correct description of Z-DNA and B-DNA is therefore conflicting.

In this work, we propose a modification that represents a reasonable compromise between Z-DNA and B-DNA molecules and provides very good results for both forms of DNA. Our OL21 α/γ modification reduces the population of non-native backbone substates in the CpG steps of Z-DNA to a relatively small number and further slightly improves the ZI/ZII equilibrium in GpC steps, providing the best description of Z-DNA backbone among current AMBER force fields.

Preliminary tests show that the OL21 α/γ modification has negligible effect on B-DNA structure. Duplex helical parameters, RMSD, BII substate percentages and backbone angle distributions are virtually unaffected by the modification. The only difference is the population of the non-canonical backbone α/γ substates, which is slightly different than that obtained with the OL15 force field, but since the total population is very small (about 1.5 %) and the substates present are known minor B-DNA substates, this does not pose any problem. Although further testing is clearly needed for a wide range of DNA structures, the improved description of non-canonical α/γ backbone substates promises better modeling of non-canonical DNA structures.

## Acknowledgements

This work was supported by Grant 17-16107S (P.J., M.Z.) from the Grant Agency of the Czech Republic.

## Supporting Information

The Supporting Information is available free of charge on the ACS Publications website at DOI:

OL21 α/γ dihedral parameters, analysis of the B-DNA parameters and their sequence dependence, B-DNA fraying and time series of selected backbone angles.

## Footnotes

^a)^ Simulation in 0.15 M KCl and SPC/E water, with original phosphate vdW parameters and flat well restraints at both ends to avoid bias from occasional fraying.

